# *Lactococcus* G423 Ameliorates the Growth Performance of Broilers by Modulation of Gut Microbiota- Metabolites

**DOI:** 10.1101/2023.03.21.533656

**Authors:** Mi Wang, Wei Ma, Chunqiang Wang, Desheng Li, Yuan Wang

## Abstract

This study aimed to explore whether *Lactococcus* G423 could ameliorate growth performance of broilers by modulation of gut microbiota-metabolites based on the 16S ribosomal RNA (rRNA) and liquid chromatography-mass spectrometry (LC-MS). A total of 640 one-day-old AA broilers were randomly divided into 4 groups (Control (CON), Lac_L, Lac_H, and ABX). Average daily gain (ADG), average daily feed intake (ADFI), and feed conversion ratio (FCR) were calculated on the 42nd day. The ileum content was harvested and immediately frozen in liquid nitrogen for 16S rRNA and LC-MS analyses. Then, the results of 16S rRNA analysis were confirmed by quantitative polymerase chain reaction (qPCR). Compared with the CON group, ADG significantly increased in the Lac_H group (*P*<0.05), and survival rate significantly decreased in the Lac_H, Lac_H, and ABX groups (all *P*<0.05). A significant difference in microbial diversity was found among the four groups. Compared with the CON group, the abundance rates of *Firmicutes and Lactobacillus* in the Lac_H group were significantly risen (*P<0.05*). The global and overview maps and membrane transport in the Lac_L, Lac_H, and ABX groups significantly changed versus those in the CON group (*P<0.05*). The results of LC-MS demonstrated that *Lactococcus* could significantly improve the levels of some metabolites (6-hydroxy-5-methoxyindole glucuronide, 9,10-DiHOME, carbamazepine-O-quinone, N-Acetyl-L-phenylalanine, and kynurenine), and these metabolites were involved in 5 metabolic pathways. Among them, the pathways of linoleic acid metabolism, phenylalanine metabolism, and pentose and glucuronate interconversions significantly changed (*P<0.05*). *Lactococcus* improved wight and survival rate of broilers through the gut microbiota, regulating the pathways of amino acid metabolism, lipid metabolism, bile acid metabolism, and carbohydrate metabolism. However, antibiotics may negatively influence the gut microbiota.

**IMPORTANCE:** Improvements in the growth rate of broiler chickens can be achieved through dietary manipulation of the naturally occurring bacterial populations while mitigating the withdrawal of antibiotic growth promoters. *Lactococcus* is industrially crucial *lactic acid bacteria*, can be incorporated into the diets of chickens to improve their growth performance. This study investigated the key mechanisms behind this progression and pinpointed *Lactococcus* improved wight and survival rate of broilers through the gut microbiota, regulating the pathways of amino acid metabolism, lipid metabolism, bile acid metabolism, and carbohydrate metabolism.

## 1. Introduction

Intestinal microbes and host are bioactive communities, forming the junction between animals and their nutritional environment (1). Thus, microbiota may affect the physiology and metabolism of host, and certain healthy bacteria in the microbiome may improve gut health (2-4). Intestinal microbes have noticeably attracted researchers’ attention recently. Over the past two decades, same studies revealed that antibiotics can alter the likely benefit of the host-microbiota interaction or relationship by regulating the microbiota (5). In the poultry industry, antibiotics have been widely used (6). However, it is essential to pay further attention to antibiotic resistance (7, 8), and long-term use of these antibiotics could cause antibiotic residues remaining in animals, in which they seriously threaten human health (9). As is well known that the basic function of probiotics is to reduce gut-related diseases by regulating and improving the intestinal microbial balance in humans (10, 11). Recently, probiotics have been found to benefit not only human health, but also animal health (12). Several studies demonstrated that beneficial effects of probiotics for the host included suppression of growth of pathogens, modulation of the immune system, improvement of nutrient metabolism, and modification of the composition of the intestinal microbiota (13-16). Especially, an appropriate amount of *lactic acid bacteria* (LAB) can regulate the microflora in the gut (17). *Lactococcus* is industrially crucial *LAB* used to produce lactic acid, pickled vegetables, buttermilk, cheese, and several types of dairy foods and drinks. In addition, they are utilized as probiotics in specific formulations. *Lactococcus* can modulate intestinal microbiota of animals (18, 19). However, they have rarely been used as probiotics versus other *LAB* genera. The ribosomal RNA (16S) rRNA gene possesses the advantage of exploring the composition of the gut microbiota of chickens (20), broiler chickens (21), Dagu chickens (22), and naked neck chickens (23). Liquid chromatography-mass spectrometry (LC-MS) has solid analytical capability, and it can detect the association of bacteria and metabolites with high resolution and accuracy (24). Moreover, correlation analysis between microorganisms and metabolites was performed. This was of great significance in revealing the contribution of bacteria to the formation of metabolites in *Lactococcus*. Therefore, the present study aimed to explore whether *Lactococcus* G423 could ameliorate growth performance of broilers by modulation of gut microbiota-metabolites based on the 16S rRNA and LC-MS.

## 2. Materials and Methods

### 2.1. Birds, diets, and experimental design

Totally, 640 one-day-old AA broilers (Shu-ya Poultry Co., Ltd., Tieling, China) were randomly assigned into four experimental groups, and each group included 160 birds (four replicates of 40 birds). Birds were raised in stainless steel cages (400×450×1500 mm^3^) in a controlled room for 42 days. The temperature of room was gradually reduced by 3.0-3.5 °C weekly until achieving a thermo-neutral zone ranged from 21 to 26 °C by the end of the 3rd week. The experimental diets were based on corn and soybean meal. Four dietary regimes were provided as follows: control group (basal diet, CON group), Lac-L and Lac-H groups (basal diet supplemented with 50 and 100 g/kg *Lactococcus*, respectively), and ABX group (basal diet supplemented with 50 g/kg narasin). The basal diet was divided into two phases: the starter phase from 1 to 21 days and the growth phase from 21 to 42 days. The basal diet was formulated to meet the nutritional requirements according to the Chinese Broiler Feeding Standards (NY/T33-2004) (**Table 1**). Broiler performance in terms of average daily gain (**ADG**), average daily feed intake (**ADFI**), survival rate, and feed conversion ratio (**FCR**) were weekly recorded, in which ADG, ADFI, and FCR were calculated and presented for 6-week experimental period.

**Table 1.**
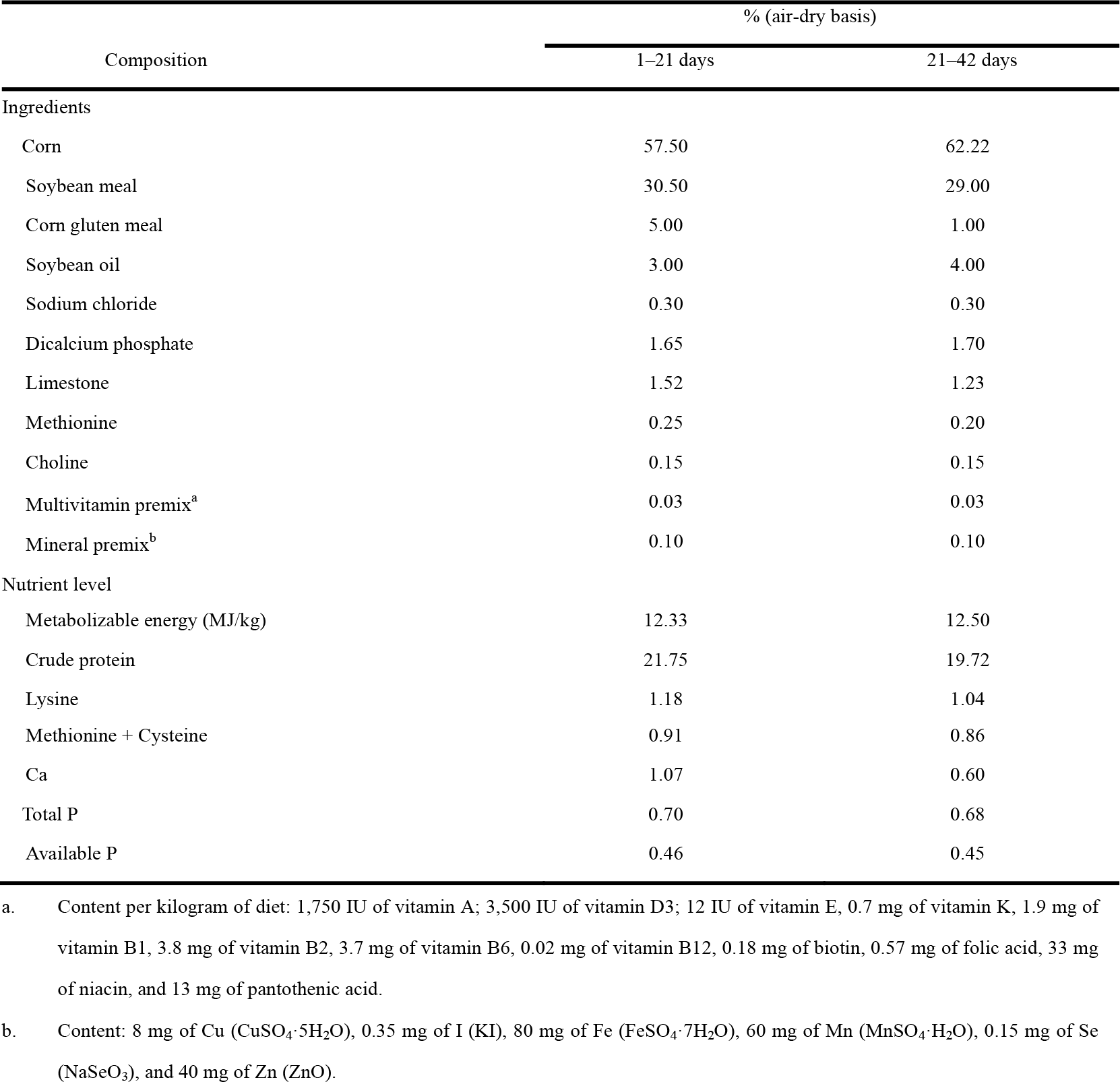
Calculated composition of basal diets and nutrient levels.

| Composition | % (air-dry basis) |  |
| --- | --- | --- |
|  | 1–21 days | 21–42 days |
| Ingredients |  |  |
| Corn | 57.50 | 62.22 |
| Soybean meal | 30.50 | 29.00 |
| Corn gluten meal | 5.00 | 1.00 |
| Soybean oil | 3.00 | 4.00 |
| Sodium chloride | 0.30 | 0.30 |
| Dicalcium phosphate | 1.65 | 1.70 |
| Limestone | 1.52 | 1.23 |
| Methionine | 0.25 | 0.20 |
| Choline | 0.15 | 0.15 |
| Multivitamin premix<sup>a</sup> | 0.03 | 0.03 |
| Mineral premix<sup>b</sup> | 0.10 | 0.10 |
| Nutrient level |  |  |
| Metabolizable energy (MJ/kg) | 12.33 | 12.50 |
| Crude protein | 21.75 | 19.72 |
| Lysine | 1.18 | 1.04 |
| Methionine + Cysteine | 0.91 | 0.86 |
| Ca | 1.07 | 0.60 |
| Total P | 0.70 | 0.68 |
| Available P | 0.46 | 0.45 |

a. Content per kilogram of diet: 1,750 IU of vitamin A; 3,500 IU of vitamin D3; 12 IU of vitamin E, 0.7 mg of vitamin K, 1.9 mg of vitamin B1, 3.8 mg of vitamin B2, 3.7 mg of vitamin B6, 0.02 mg of vitamin B12, 0.18 mg of biotin, 0.57 mg of folic acid, 33 mg of niacin, and 13 mg of pantothenic acid.
b. Content: 8 mg of Cu (CuSO<sub>4</sub>·5H<sub>2</sub>O), 0.35 mg of I (KI), 80 mg of Fe (FeSO<sub>4</sub>·7H<sub>2</sub>O), 60 mg of Mn (MnSO<sub>4</sub>·H<sub>2</sub>O), 0.15 mg of Se (NaSeO<sub>3</sub>), and 40 mg of Zn (ZnO).

### 2.2 Illumina MiSeq sequencing for detection of intestinal microbial diversity

Four ileal samples per group were randomly selected for the analysis of intestinal flora. The polymerase chain reaction (PCR) amplification of the hypervariable region V3-V4 of the 16S rRNA gene was performed with the universal primers set338 F (5′-ACTCCTACGGGAGGCAGCAG-3′) and 806R (5′-GGACTACHVGGGTWTC TAAT-3′). The quality and concentration of DNA were determined by 1.0% agarose gel electrophoresis and a NanoDrop® ND-2000 spectrophotometer (Thermo Fisher Scientific Inc., Waltham, MA, USA) and kept at -80 ℃ for further experiment. All samples were amplified in triplicate. The PCR products were extracted from 2% agarose gel and purified using the AxyPrep DNA Gel Extraction kit (Axygen Biosciences, Union City, CA, USA) according to the manufacturer’s instructions and quantified using Quantus™ Fluorometer (Promega, Madison, WI, USA). The Illumina MiSeq platform (Illumina Inc., San Diego, CA, USA) was used for paired-end sequencing (2 × 300) of the PCR products. The raw sequence reads were quality-filtered and merged by FLASH before open-reference operational taxonomic units (OTUs) picking via UPARSE and taxonomy classification through the SILVA 16S rRNA database.

#### 2.2.2 Quantitative PCR (qPCR)

*Lactobacillus* and *Firmicutes* were detected by qPCR. Four ileum contents from broiler were collected. The primers used for qPCR are presented in Table 2. The conditions of PCR reaction were summarized as follows: (1) at 95 ℃ for 5 min; (2) a: at 95 ℃ for 30 s; b: at 60 ℃ for 30 s; c: at 72 ℃ for 1 min, including a total of 35 cycles; (3) a: at 95 ℃ for 30 s; b: at 55 ℃ for 30 s; c: at 72 ℃ for 1 min. The △Ct was calculated as follows (corrected sample) = mean value of target gene–mean value of internal reference gene (△△ Ct = △Ct–mean value of control group).

**Table 2.**
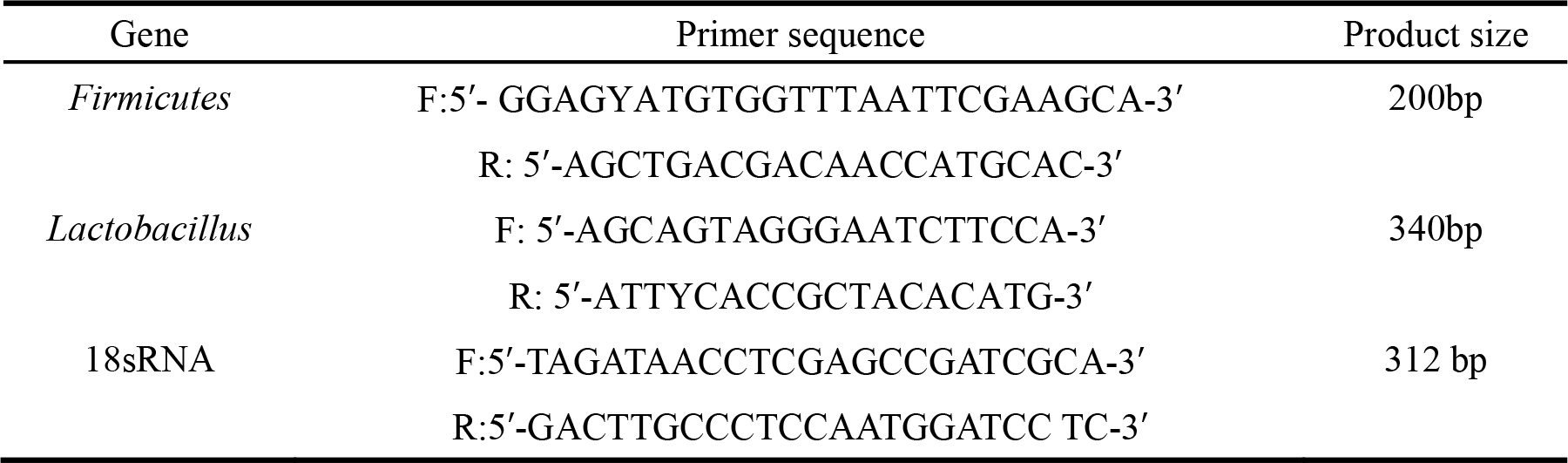
The primers used for Qpcr.

### 2. 3 LC-MS analysis

The LC-MS analysis of ileal contents was conducted on a Thermo UHPLC-Q Exactive HF-X system equipped with an ACQUITY HSS T3 column (100 mm × 2.1 mm i.d., 1.8 μm; Waters Corp., Milford, MA, USA) at Majorbio Bio-Pharm Technology Co., Ltd. (Shanghai, China). The mass spectrometric data were collected using a Thermo UHPLC-Q Exactive HF-X Mass spectrometer equipped with an electrospray ionization (ESI) source operating in positive and negative modes. The pretreatment of LC-MS raw data was performed by Progenesis QI software (Waters Corp.), and a three-dimensional (3D) data matrix in CSV format was exported. This 3D matrix included the following information: sample information, metabolite name, and mass spectral response intensity. Internal standard peaks and any known false positive peaks (including noise, column bleed, and derivatized reagent peaks) were removed from the data matrix, deredundant and peak pooled. Moreover, the metabolites were identified by searching in the following databases: Human Metabolome Database (HMDB) (http://www.hmdb.ca/), Metlin (https://metlin.scripps.edu/), and Majorbio (https://cloud.majorbio.com).

### 2.4 Statistical analysis

Between-group statistical differences were compared using one-way analysis of variance (ANOVA), followed by post-hoc multiple comparisons using the Fisher’s least significant difference (LSD) t-test. The experimental data were presented as the mean ± standard error of the mean (SEM), analyzed using SPSS 20.0 software (IBM, Armonk, NY, USA), and *P*<0.05 was considered statistically significant. The 16S rRNA genes of gut microbiota were analyzed using an online platform (https://cloud.Majorbio.com). The multivariate statistical analysis was performed using “ropls” (Version 1.6.2) R package from Bioconductor on Majorbio Cloud Platform (https://cloud.majorbio.com).

## 3. Results

### 3.1 Growth performance

There were no significant changes in FCR and ADFI (*P*>0.05) among Lac_H, Lac_L, and ABX groups, however, ADG significantly increased in the Lac_H group (*P*<0.05), and survival rate significantly decreased in Lac_H, Lac_L, and ABX groups (Table 3).

**Table 3.** Effects of *Lactococcus* on the growth performance in broilers.

|  | 42 d |  |  |  |
| --- | --- | --- | --- | --- |
|  | CON | Lac_L | Lac_H | ABX |
| ADG (g) | 59.17±0.87<sup>a</sup> | 60.30±1.07<sup>a</sup> | 62.80±0.96<sup>b</sup> | 60.08±0.75<sup>a</sup> |
| ADFI (g) | 88.99±0.77<sup>a</sup> | 88.28±0.75<sup>a</sup> | 91.59±3.14<sup>a</sup> | 88.27±0.95<sup>a</sup> |
| Survival rate (%) | 91.4±3.66<sup>a</sup> | 96.8±0.92<sup>b</sup> | 97.8±1.03<sup>b</sup> | 95.9±1.15<sup>b</sup> |
| FCR | 1.50±0.028<sup>a</sup> | 1.47±0.018<sup>a</sup> | 1.46±0.022<sup>a</sup> | 1.47±0.014<sup>a</sup> |

<sup>1</sup>ADFI = average daily feed intake; <sup>2</sup>ADG = average daily gain; <sup>3</sup>FCR = feed conversion ratio. <sup>4</sup>In the same row, values with different superscripts represent significant differences (*P*<0.05). Values are expressed as mean ± SEM (n=4 for all groups).

### 3.2 Intestinal microflora

To characterize the intestinal microbiota composition of broilers in the four groups, 16S rRNA gene sequence analysis was performed. A total of 744461 clean sequences were obtained. With the sequence similarity of 97%, 702 OTUs were obtained. The average good’ s coverage for samples was higher than 99%, indicating that the majority of the microbial species were identified and sequencing depth was also adequate for the robust sequence analysis.

As illustrated in **Table 4**, alpha diversity analysis of gut microbiota showed that compared with the CON group, the Chao and Ace indices in the Lac_H, Lac_L, and ABX groups significantly increased (P < 0.05), however, the Simpson index exhibited an opposite trend. The Simpson index in the Lac_L, Lac_H, and ABX groups was significantly reduced compared with that in the CON group *(P < 0.05*). In addition, the Sob index in the Lac_H and ABX groups was significantly higher than that in the CON group (*P < 0.05*). The Shannon index in the Lac_L and Lac_H groups was significantly elevated compared with that in the CON group (P < 0.05). The Coverage index in the Lac_L and ABX groups significantly increased compared with that in the CON group (*P < 0.05*). The effects of *Lactococcus* on the diversity and richness of intestinal microbiota community in crucian carp were evaluated based on alpha diversity (**Table 4**).

**Table 4.** Alpha diversity of intestinal microbiota based on OTU levels.

| Index | CON | Lac_L | Lac_H | ABX |
| --- | --- | --- | --- | --- |
| Coverage | 0.99986±0.0001<sup>a</sup> | 0.99914±0.000134<sup>b</sup> | 0.99934±0.000433a | 0.99929±0.000269<sup>b</sup> |
| Sobs | 15±4<sup>a</sup> | 120±65.483<sup>a</sup> | 230.33±69.601<sup>b</sup> | 177.33±82.008<sup>b</sup> |
| Shannon | 0.78077±0.118122<sup>a</sup> | 1.2905±0.20873<sup>b</sup> | 2.3078±0.84541<sup>b</sup> | 2.074±1.082<sup>ab</sup> |
| Simpson | 0.60533±0.064049<sup>a</sup> | 0.40857±0.076413<sup>b</sup> | 0.20462±0.08412<sup>c</sup> | 0.24308±0.13222<sup>bc</sup> |
| Ace | 44.601±28.667<sup>a</sup> | 173.16±50.96<sup>b</sup> | 253.89±75.984<sup>b</sup> | 197.65±75.663<sup>b</sup> |
| Chao | 32±22.068<sup>a</sup> | 154.38±68.299<sup>b</sup> | 255.83±79.422<sup>b</sup> | 201.12±76.12<sup>b</sup> |

In the same row, values with different superscripts indicate significant differences ( $P<0.05$ ). Values are expressed as mean ± SEM (n=4 for all groups).

Principal coordinate analysis (**PCoA**) based on OUT abundance showed that points in the Lac_H and ABX groups were scattered in the right, which indicated that the microbial structure in the Lac_H and ABX groups had undergone a tremendous change versus that in the CON group. In the Lac_L and CON groups, points were clustered separate from each other in the left, which showed that the low dose of *Lactobacillus* could change the structure of gut microbiota (Fig. 1A). At the phylum level, *Firmicutes, Proteobacteria*, and *Bacteroidetes* were the top species identified in all samples (Fig. 1B). At the genus level, compared with those in the CON group, the abundance of *Lactobacillus* significantly increased, whereas that of *Bacteroides* significantly decreased in the Lac_L and Lac_H groups (Fig. 1C). As illustrated in Fig. 1D, qPCR showed that the proportion of *Lactobacillus* in the Lac_L and Lac_H groups was significantly elevated compared with that in the CON group (*P<0.05*). Additionally, proportion of *Firmicutes* was significantly risen after treating with Lac_H (*P<0.05*). The different effects of Lac_L and Lac_H on microbiota might justify their different biochemical parameters. Subsequent linear discriminant analysis effect size (**LEfSe**) revealed substantial differences in *Lactobacillus_ salivarius* and *Lactobacillus_ johnsonii* in the Lac_H and ABX groups (Fig. 1E).

**Fig.1.**
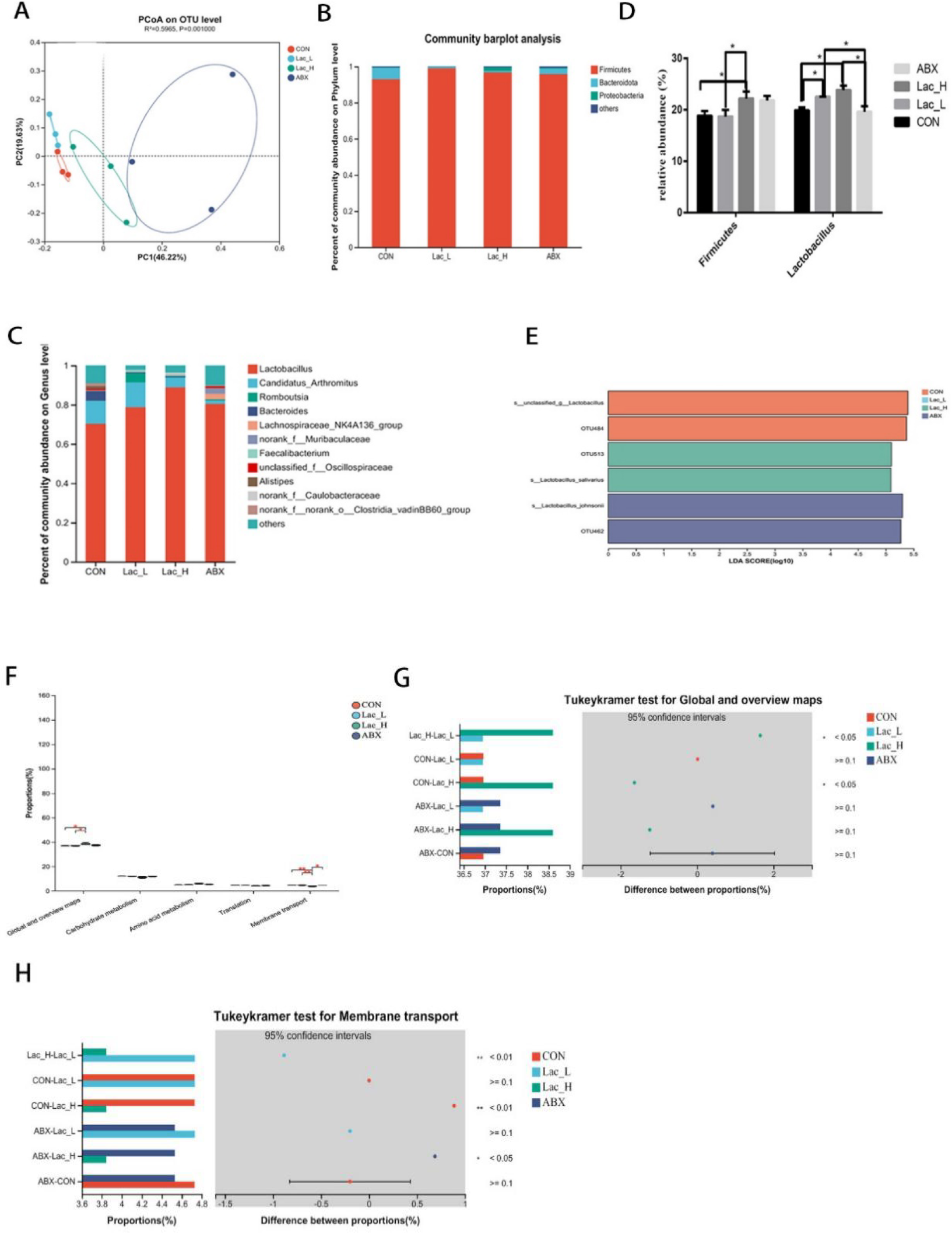
Effects of *Lactococci* on gut microbiota of broilers. Principal coordinate analysis (PCoA) based on the weighted UniFrac distance (A); Column chart of community difference at the phylum level (B) and the genus level (C); Relative abundance of discriminative gut microbiota at the genus level (D); LEfSe analysis (E); KEGG pathway analysis (F-H); * and ** represent P < 0.05 and P < 0.01, respectively.

The function of the ileum microbiome was predicted using the Phylogenetic investigation of communities by reconstruction of unobserved species 2 (**PICRUSt2**). Then, the Kyoto Encyclopedia of Genes and Genomes **(KEGG**) pathway analysis was used to divide the predicted metabolic pathways into six functional groups. The microbial communities in the CON, Lac_L, Lac_H, and ABX groups were mainly related to metabolism, genetic information processing, cellular processes, environmental information processing, human diseases, and organic systems. Their main functions were concentrated in the metabolism of amino acids, carbohydrate, vitamins, terpenoids, polyketides, and lipids (Table 4).

As displayed in Fig. 1F, the global and overview maps and membrane transport in the Lac_L, Lac_H, and ABX groups significantly changed compared with those in the CON group (*P <0.05*). Functional predictions of differences in the mean relative abundance among groups are shown in Fig. 1G-H and Table 5.

**Table 5.**
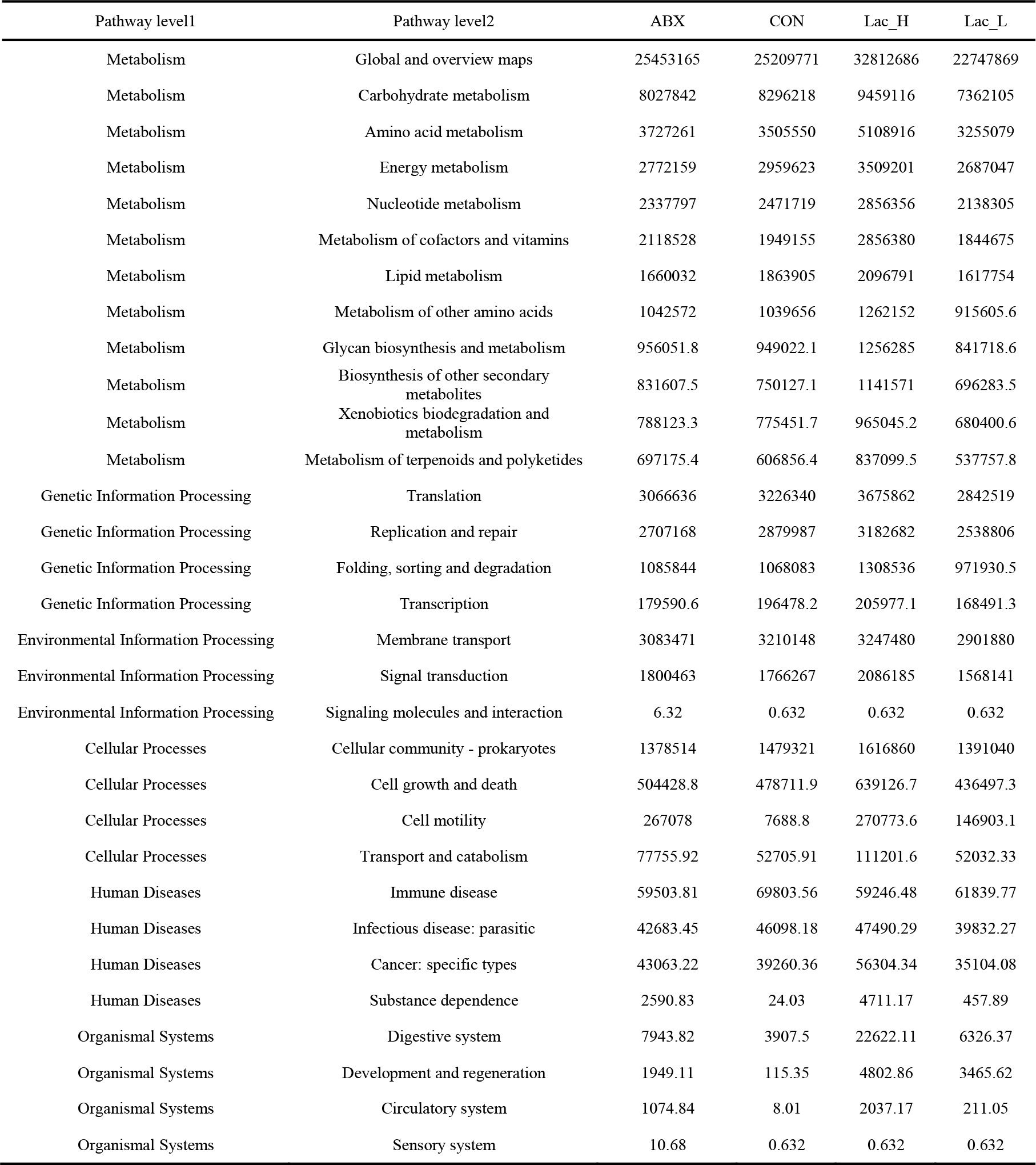
Functional prediction of colonic microbiota in broilers.

### 3.3 Intestinal Metabolites

A total of 6612 and 5851 metabolites in ileal contents were determined in positive and negative ion modes, respectively, using LC-MS-based nontargeted metabolomics. A total of 228 metabolites were identified and named based on the HMDB and KEGG databases. Furthermore, orthogonal projection to latent structures-discriminant analysis (OPLS-DA) was employed to select the most predictive and discriminative features to assist classify cation. The loading plot showed a clear separation in metabolites between the Lac_H and CON groups (Figure 2A). The results revealed that the metabolite of broiler significantly changed after treating with *Lactococcus*. Then, the heat map tree of cluster analysis of metabolites (Figure 2B) was constructed, which visualized 50 significantly different metabolites. Overall, there were significant differences in metabolites between the CON and Lac_H groups.

**Fig. 2.**
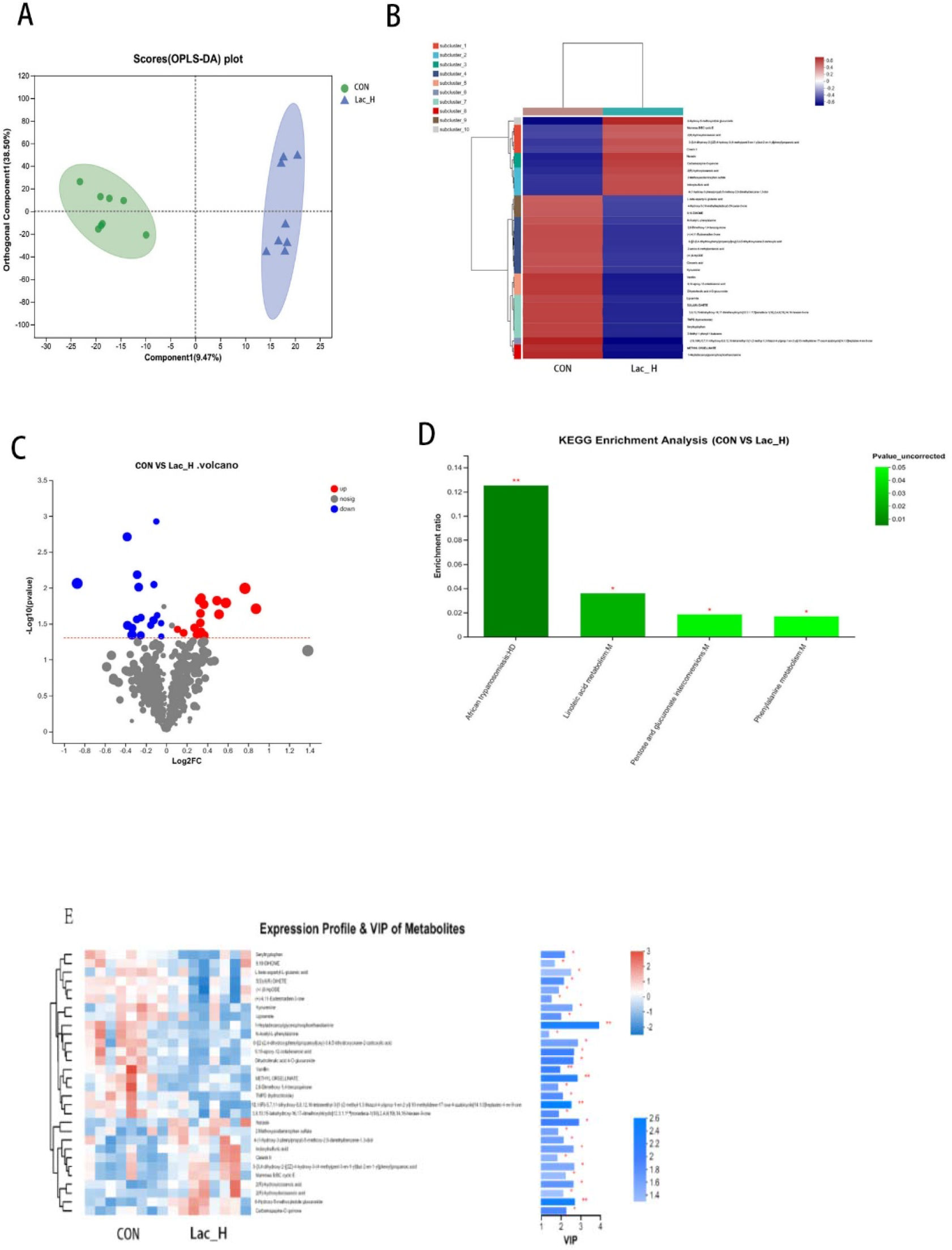
Effects of Lac_H on ilea metabolites of broilers. Multivariate statistical analysis of blank control group and Lac_H group (A); Tthe heat map of cluster analysis of metabolites (B); Volcanic diagram of differentially expressed metabolites (C); Variable importance in projection (VIP) scores of the CON group vs. Lac_H group (D). Bubble diagram of metabolic pathway enrichment analysis (E); *, **, and *** represent P < 0.05, P < 0.01, and P < 0.001, respectively.

The levels of several metabolites, such as 6-hydroxy-5-methoxyindole glucuronide, 3-{3,4-dihydroxy-2-[(2Z)-4-hydroxy-3-(4-methylpent-3-en-1-yl)but-2-en-1-yl]phenyl}propanoic acid, indoxylsulfuric acid, Cinerin II, carbamazepine-O-quinone, and 4-(1-hydroxy-3-phenylpropyl)-5-methoxy-2,6-dimethylbenz ene-1,3-diol were upregulated in the Lac_H group, while the levels of 9,10-DiHOME, seryltryptophan, seryltryptophan dihydroferulic acid 4-O-glucuronide, 2,6-dimethoxy-1,4-benzoquinone, and kynurenine were downregulated (Figure 2C). Metabolites discriminated among different groups were screened using the variable importance in projection (VIP) scores obtained from the OPLS-DA model, and the ileal contents of metabolic profiles were determined. The metabolites were statistically significant if VIP score ≥ 1 and *P* < 0.05, and *P* was calculated by the *t*-test. Metabolites with VIP score > 1.0 and P < 0.05 were considered to be significantly influenced by the Lac_H. Thirty significantly affected metabolites were identified in the CON and Lac_H groups, respectively; and the top 30 metabolites with the highest VIP scores are presented in Fig. 2D.

Metabolic pathway enrichment analysis was performed based on the KEGG database for the differential metabolites between the CON and Lac_H groups, and the metabolic pathway with *P <* 0.05 was significantly enriched for the differential metabolites, which including Bile secretion, Linoleic acid metabolism, Drug metabolism - cytochrome P450, Phenylalanine metabolism, Tryptophan metabolism, matches metabolites. *Lactococcus* could significantly improve the levels of certain metabolites (6-hydroxy-5-methoxyindole glucuronide, 9,10-DiHOME, carbamazepine-O-quinone, N-acetyl-L-phenylalanine, and kynurenine), and these metabolites were involved in 5 metabolic pathways (Table 6). Among them, the pathways of linoleic acid metabolism, phenylalanine metabolism, and pentose and glucuronate interconversions significantly varied (*P <* 0.05) (Fig. 2E).

**Table 6.**
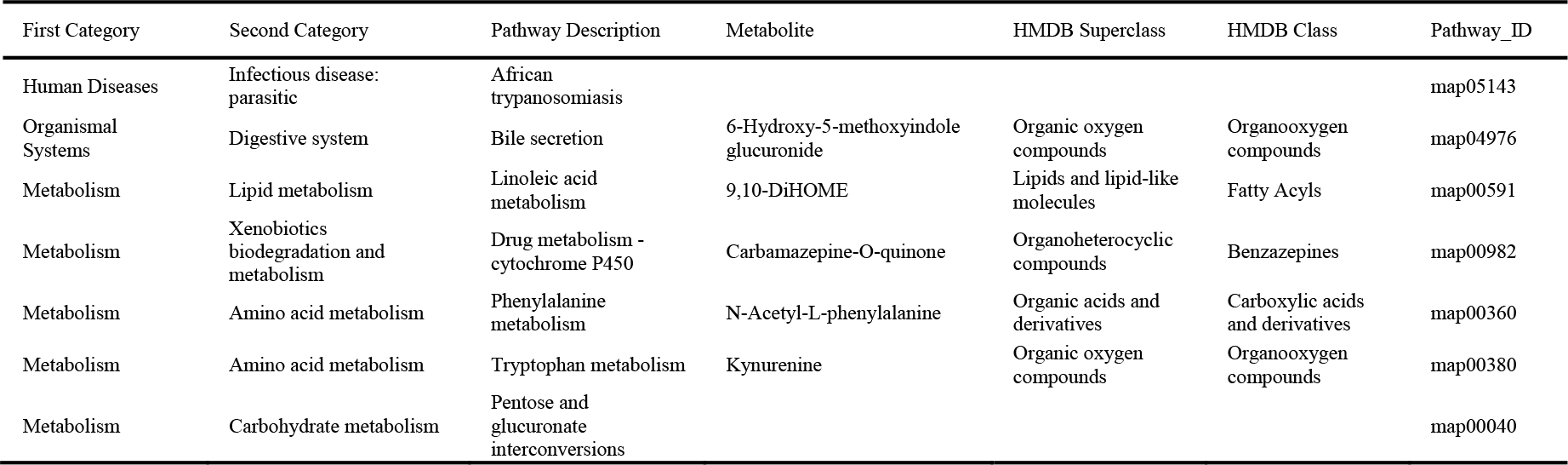
The effects of *Lactococcus* on the changes of intestinal metabolic pathway in broilers.

### 3.3 Correlation analysis between metabolites and intestinal microbiota

The variations in the intestinal microbiota could be related to the metabolic phenotype. As show in Fig. 3, correlation analysis was performed between 34 different metabolites and 44 bacteria with significantly different relative abundances at the genus level. There was a significant correlation between 2(R)-hydroxyicosanoic acid, 2(R)-hydroxydocosanoic acid, L-beta-aspartyl-L-glutamic acid, 9,10-DiHOME, TMPD (hydrochloride), kynurenine, 6-hydroxy-5-methoxyindole glucuronide, seryltryptophan, N-acetyl-L-phenylalanine and *Parabacteroides, Romboutsia, Sellimonas, Subdoligranulum, Turicibacter, Tuzzerella, Bacteroides, Lachnospiraceae, Butyricicoccus, Candidatus_Arthromitus, Eisenbergiella, Escherichia-Shigella, Faecalibacterium, Alistipes, Marvinbryantia, Monoglobus, Negativibacillus* (all P < 0.05).

**Fig.3.**
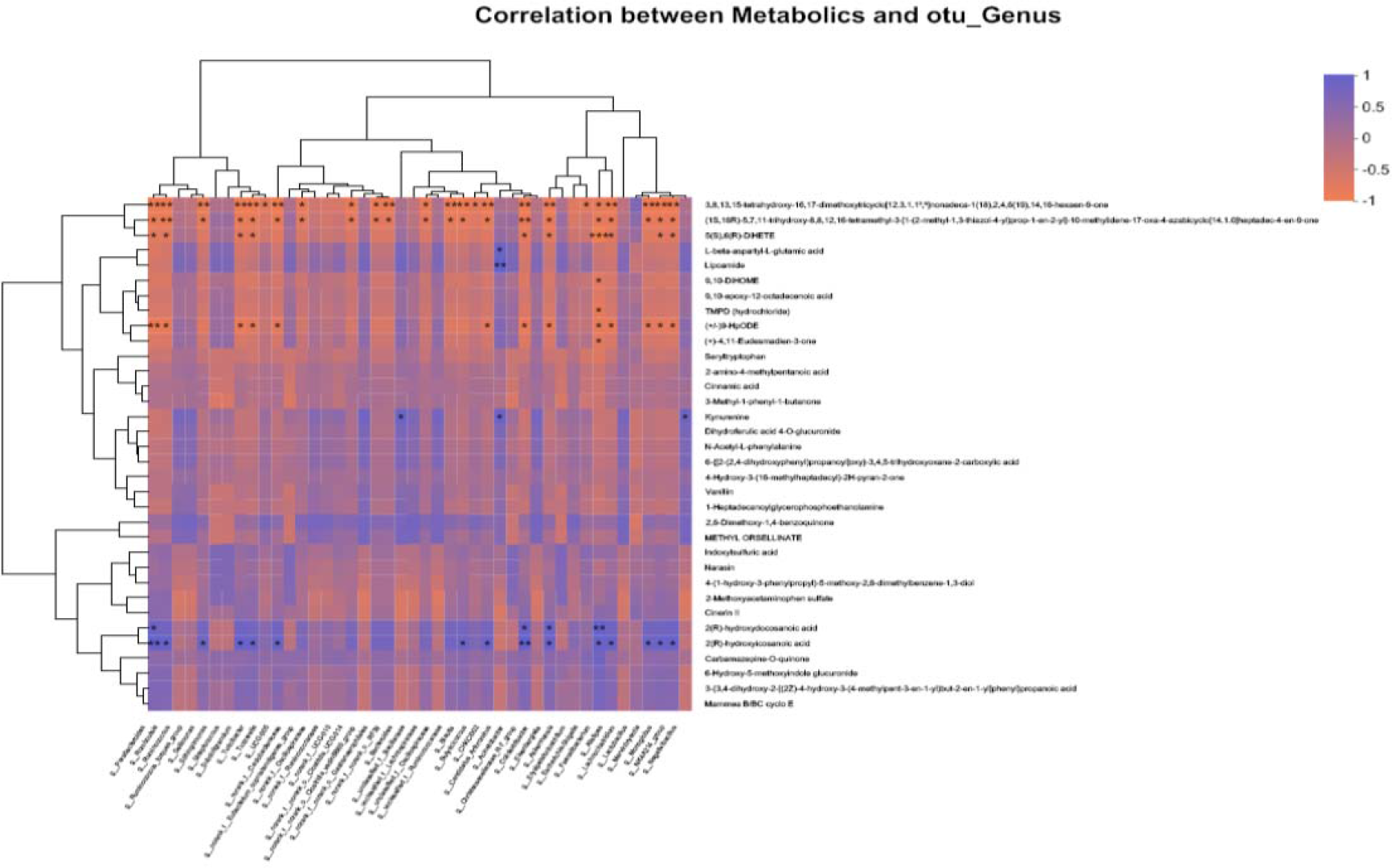
Correlation analysis of “metabolites-intestinal flora” in broilers. Note: Horizontal coordinates indicate metabolites and vertical coordinates indicate gut microbiota; R values are shown in different colors in the graph, in which red indicates positive correlation and blue indicates negative correlation; *, **, and *** represent P < 0.05, P < 0.01, and P < 0.001 respectively.

## 4. Discussion

The results of the present study showed that the gut microbiota and the ileum contents of metabolites were significantly correlated, and the metabolites might be considered as mediators in the association between the intestinal microbiota and growth performance. *Lactococcus* could ameliorate the growth performance of broilers by integrating the microbiome and metabolome data.

The diversity and relative abundance of intestinal microbes play an important role in the health of host by participating in metabolism and immunomodulation (25). The findings of the present study suggested that *Lactococcus* could significantly increase ADG and improve survival rate in broilers, which were similar to previously reported results (26, 27). Although the mechanism underlying the growth-promoting action in broilers has still remained elusive, a main influential factor may be an increase in nutrient digestibility. The gut microbiota community is consisted of diverse types of microbes. In the present study, it was found that *Lac* and ABX altered microbiome diversity in ileum of broilers, and changed the relative abundance rates of *Firmicutes, Bacteroidetes, Proteobacteria*, and other species. At the phylum level, *Firmicutes, Bacteroidetes*, and *Proteobacteria* were the most common phyla in the poultry intestinal samples, which is consistent with previous findings (28-30). This study indicated that *Firmicutes* were the dominant phylum (> 50%) in broilers, and similar results have been previously reported (31,32) Moreover, this study revealed that the abundance rates of *Firmicutes* were relatively higher in the Lac_L and Lac_H groups compared with those in the CON group (*P<0.05*). *Firmicutes* have been considered beneficial to the host (33). Turnbaugh et al. reported that *Firmicutes* were associated with lipid metabolites (34). The phylum *Bacteroidetes* have influences on dissolving proteins, lipids, and carbohydrates (35, 36). Some studies demonstrated that the *Firmicutes/Bacteroidetes* ratio and the growth performance was positively correlated, and this ratio could be indicative of the status of the intestinal bacteria (37,38). The *Firmicutes/Bacteroidetes* ratio was a higher in obese children (39). The results of the present study revealed that *Firmicutes/Bacteroidetes* ratio was relatively higher in the Lac_L, Lac_H, and ABX groups compared with that in the CON group. *Lactococcus* significantly increased ADG by changing *Firmicutes/Bacteroidetes* ratio. Additionally, the levels of *Proteobacteria* phylum, including some pathogens, such as *Escherichia, Salmonella, Helicobacter*, and *Vibrio*, were slightly lower in the Lac_L group than those in the CON group, indicating that *Lactococcus* decreased the abundance of opportunistic pathogens. However, at the genus level, *Lactobacillus, Candidatus_Arthromitus, Romboutsia*, and *Bacteroides* were identified as the dominant species in the ileum microbiome. *Lactobacillus*, belonging to the phylum of *Firmicutes*, level in the Lac_L and Lac_H groups was markedly higher than that in the CON group (*P<0.05*), and the abundance of *Lactobacillus* in the Lac_ H group reached the highest rate.

*Lactobacillus* is involved in digestive and metabolic processes, as well as in the regulation of local and systemic immune response (40). And *Lactobacillus* promotes intestinal health in humans and animals by producing a variety short-chain fatty acids. Moreover, *Lactobacillus* promotes the gut health of humans and animals via production of various short-chain fatty acids. *Lactobacillus* participates in gut microbiota, growth and nutrient utilization, immune system, and mineral metabolism in animals (41). *Lactobacillus* has a positive effect on the growth and fat deposition in broilers (42). Therefore, *Lactococcus* increases the abundance of *Lactobacillus*, which is the primary outcome about the health of broilers. In the present study, it was revealed that the abundance of *Bacteroides* in the Lac_L, Lac_H, and ABX groups was markedly lower than that in the CON group. *Bacteroides* are also positively correlated with several lipid metabolites (43,44). Yun Huang et al. (45) reported that the genus *Bacteroides* had a negative FCR-associated correlation. The appropriate amount of *Bacteroides* might be able to degrade non-starch polysaccharides (46,47).There was a negative association between body weight and the abundance of *Bacteroidetes* (48,49). *Romboutsia* has been identified in the human gut (50), the rat gastrointestinal tract (51), and the fecal of hens (52). A novel genus *Romboutsia* was found in ileum samples of broilers in the present study. However, the abundance of *Romboutsia* was inconsistent among the four groups. Meanwhile, it was revealed that ABX altered relative abundance rates of other bacteria in ileum contents of broilers, negatively influencing the gut microbiota. It was found that *Lactococcus* could regulate lipids, proteins, and carbohydrates by gut microbiota.

The *Lactococcus*-regulated gut microbiota led to alterations in the contents of ileum metabolites. Several significantly altered metabolites were identified in the present study, such as 6-hydroxy-5-methoxyindole glucuronide, 9,10-DiHOME, carbamazepine-O-quinone, N-acetyl-L-phenylalanine, and kynurenine, which were regulated and they were involved in 5 metabolic pathways (Table 5). Among them, the bile secretion metabolism is an important metabolic pathway of glucose and lipid (53). It was revealed that 6-hydroxy-5-methoxyindole glucuronide was related to the pathway of bile acid metabolism. Bile acids are also signaling molecules and inflammatory agents that rapidly activate nuclear receptors and cellular signaling pathways, regulating lipid, glucose, and energy metabolism. Bile salts are to a large extent (>95% per cycle) absorbed in the terminal ileum, the final section of the small intestine. The bile salt hydrolase activity has been widely detected in several bacterial genera, including *Bacteroides, Clostridium, Lactobacillus, and Bifidobacteria*. In addition, bile acids have been found to affect glucose metabolism by activating FXR and TGR5 receptors, as well as intestinal flora (54). *Lactococcus* upregulated 6-hydroxy-5-methoxyindole glucuronide level, suggesting that it may have some regulatory effects on the bile acid metabolism.

Linoleic acid has shown a positive correlation with metabolic diseases (55). Previous studies have demonstrated that a lower linoleic acid content was associated with an increased risk of cardiovascular disease and type 2 diabetes mellitus, and a higher linoleic acid content was correlated with a lower risk of developing metabolic syndrome (56-58). Linoleic acid is metabolized to produce 9,10-dihydroxy-12-octadecenoic acid (9,10-DiHOME), which may induce oxidative stress by activating nuclear factor-κB (NF-*k*B) and activator protein-1 (AP-1) transcription factors that mediate inflammation (59). *Propionibacterium acnes* and *Lactobacillus plantarum* have been reported to convert linoleic acid into conjugated linoleic acid (60). It has been proved that conjugated fatty acids have many potential physiological properties including anti-carcinogenic, anti-obesity, anti-cardiovascular and anti-diabetic activities (60). In the present study, *Lactococcus* downregulated 9,10-DiHOME level, suggesting that it may have some regulatory effects on the linoleic acid metabolism.

In addition, N-acetyl-L-phenylalanine is a product of enzyme phenylalanine N-acetyltransferase, which is found in the phenylalanine metabolism pathway (61). Kynurenine is the key metabolite in the tryptophan metabolic pathway, which has been specifically studied in neural and immune systems (62). Tryptophan may be degraded into kynurenines by *pseudomonas aeruginosa* through the kynurenine pathway (63,64). It is noteworthy that phenylalanine and tryptophan are 2 of the 20 amino acids used for synthesizing proteins with the unique characteristic. It was reported that peptides with a high rate of tryptophan could ameliorate anxiety/depression-like behaviors via the kynurenine pathway (65). Mammalian indoleamine 2,3-dioxygenases regulate infection and drive immune tolerance by tryptophan deprivation and the generation of active metabolites, such as kynurenines (66). In the present study, *Lactococcus* downregulated kynurenine level, suggesting that it may have some regulatory effects on amino acid metabolism. Notably, correlation analysis between diversity and metabolites of ileum in broilers indicated that bacteria, such as *Lactobacillus, Candidatus_Arthromitus, Romboutsia*, and *Bacteroides* were associated with 9,10-DiHOME, kynurenine, 6-hydroxy-5-methoxyindole glucuronide, and N-acetyl-L-phenylalanine in the CON group versus the Lac_H group.

## Conclusions

In conclusion, *Lactococcus* improved weight and survival rate of broilers through the gut microbiota, regulating the pathways of amino acid metabolism, lipid metabolism, bile acid metabolism, and carbohydrate metabolism. However, antibiotics may negatively affect the gut microbiota. The results revealed that *Lactobacillus* had more noticeable health benefits compared with antibiotics. Thus, the above-mentioned *Lactococcus* strains can be utilized as a new probiotic combination for humans and animals.

## Data Availability

The data used to support the findings of this study are available from the corresponding author upon reasonable request.

## Conflict of interest

The authors declare that there is no conflict of interest.

## Authors’ Contributions

Mi Wang participated in the design of the study and critically revised the manuscript. Wei Ma and Chunqiang Wang provided technical support for the experiments. Yuan Wang performed the experiments and participated in the statistical analysis. Desheng Li contributed to the revision of the manuscript. All the authors read and approved the submitted version of the manuscript.

## Acknowledgments

This study was supported by the Crosswise Tasks (Grant No. 2021010).

